# MdrDB: Mutation-induced drug resistance DataBase

**DOI:** 10.1101/2022.10.20.513118

**Authors:** Ziyi Yang, Zhaofeng Ye, Jiezhong Qiu, Rongjun Feng, Danyu Li, Changyu Hsieh, Jonathan Allcock, Sheng-Yu Zhang

## Abstract

Mutation-induced drug resistance – where the efficacy of drugs is diminished by structural changes in proteins – presents a significant challenge to drug development and the clinical treatment of disease. Understanding the effects of mutation on protein-ligand binding affinities is a key step in developing more effective drugs and therapies, but as a research community we are currently hindered by the lack of a comprehensive database of relevant information. To address this issue, we have developed MdrDB, a database of information related to changes in protein-ligand affinity caused by mutations in protein structure. MdrDB combines data from seven publicly available datasets with calculated biochemical features, as well as 3D structures computed with PyMOL and AlphaFold 2.0, to form the largest database of its kind. With 3D structural information provided for all samples, MdrDB was specifically created to have the size, breadth, and complexity to be useful for practical protein mutation studies and drug resistance modeling. The database brings together wild type and mutant protein-ligand complexes, binding affinity changes upon mutation (ΔΔG), and biochemical features calculated from complexes to advance our understanding of mutation-induced drug resistance, the development of combination therapies, and the discovery of novel chemicals. In total, MdrDB contains 100,537 samples generated from 240 proteins (5,119 total PDB structures), 2,503 mutations, and 440 drugs. Of the total samples, 95,971 are based on available PDB structures, with the remaining 4,566 based on AlphaFold 2.0 predicted structures.

## 1 Introduction

Structural mutation of proteins can directly affect their folding and stability [1, 2, 3], function [4, 5], interactions with other proteins [6, 7] and binding affinity [8]. In some cases, it can result in significant perturbations or even complete abolishment of protein function, potentially leading to diseases or cancer [9, 10]. The evolutionary pressure imposed by small molecule drugs on many quickly evolving systems, including cancer cells, viruses, and bacteria, can lead to the rapid development of resistance [11, 12, 13, 14]. While novel and cheap high-throughput sequencing technologies have made it possible to identify mutations in large populations [15, 16], the significance and characteristics of any novel polymorphisms currently requires time-consuming and expensive experiments to determine [17].

Protein-ligand binding affinity data is of great value for understanding the impact of polymorphisms on disease and identifying mutations that lead to drug resistance. Convenient and broad access to such data for wild type and mutant proteins would aid our understanding of the mechanisms of mutation-induced drug resistance, increase the accuracy of extrapolations to novel mutations and systems, enable more effective computational approaches for drug resistance prediction, and facilitate the development of combination therapies and the discovery of novel chemicals.

Fortunately, a number of databases on the effects of mutation on protein-ligand binding affinity have been compiled and released in recent years. In particular, Platinum [17] was the first database to provide experimental data on changes in protein–ligand affinities upon mutation, along with three-dimensional structures; while TKI [18] contains reliable inhibitor ΔpIC50 data for 144 clinically identified mutants of the human kinase Abl. Together, these two datasets are the current gold-standards, both providing well-curated data and three-dimensional structural information. Other resources include AIMMS [19], RET [20] and KinaseMD [21], which provide kinase inhibitor resistance information. However, while the value of these datasets is significant, there remain a number of deficiencies. Platinum and TKI are restricted to known co-crystal structures and thus contain relatively few samples (approximately 1,000 and 150, respectively), which can limit their use with machine learning models. For AIMMS, RET, and KinaseMD, structural information is not provided. In addition, while all of the aforementioned databases consider single-point and multi-site substitution mutations, they do not include more varied and complex mutation types – such as deletions, insertions, and insertion-deletions (indel) – which play an important role in certain disease progressions.

To address these issues, we have developed MdrDB, a comprehensive database of information which combines data from all five of the databases mentioned above, with information from the Genomics of Drug Sensitivity in Cancer (GDSC) dataset [22] and DepMap [23], which provides a systematic measure of drug response to somatic mutations in different cancer cell lines. Further supplemented with data from supporting databases such as RCSB PDB, UniProtKB and PubChem, and 3D structures computed with PyMOL and AlphaFold 2.0 [24], MdrDB brings together wild type and mutant protein-ligand complexes, binding affinity changes upon mutation (ΔΔG), and biochemical features calculated from complexes to advance our understanding of mutation-induced drug resistance. MdrDB contains 100,537 samples, generated from 240 proteins (5,119 total PDB structures), 2,503 mutations, and 440 drugs. 95,971 samples are based on available PDB structures, and 4,566 samples are based on AlphaFold 2.0 predicted structures.

MdrDB offers three key advantages over existing publicly available protein mutation databases, and was created to have the size, breadth, and complexity to be useful for practical protein mutation studies and drug resistance modeling:

- **Database size:** With over 100,000 samples, MdrDB is, to our knowledge, the largest mutation-induced drug resistance database, integrating information from multiple sources, and covering mutations across various protein families. A database of this size and breadth enables the development of more effective data-driven models for drug resistance prediction.
- **Structure-based information:** Unlike KinaseMD and GDSC, MdrDB provides 3D structural information on all wild type and mutant proteins included in the database. Such structural information is critical for both accurate feature calculations used in computational drug design as well as the visualization of the effects of mutation. Based on these structures, a total of 146 biochemical features were calculated to provide data for drug resistance modeling.
- **Diverse protein mutations:** In addition to single-point and multi-site substitution mutations considered in existing databases, MdrDB contains a variety of complex mutation types such as deletions, insertions, and insertion-deletions, which can play an important role in some disease progressions.

## 2 DATA COLLECTION AND PROCESSING

The MdrDB database construction pipeline is shown in Figure 1. Data preparation includes four steps: (1) Data collection from the seven publicly available data sets. (2) Data preprocessing and integration of information from other databases, so that each sample contains information on the protein, drug, mutation and ΔΔG. (3) 3D structure generation for the wild type and mutant protein-drug complexes. (4) Calculation of biochemical features based on the complex structures.

**Figure 1:**
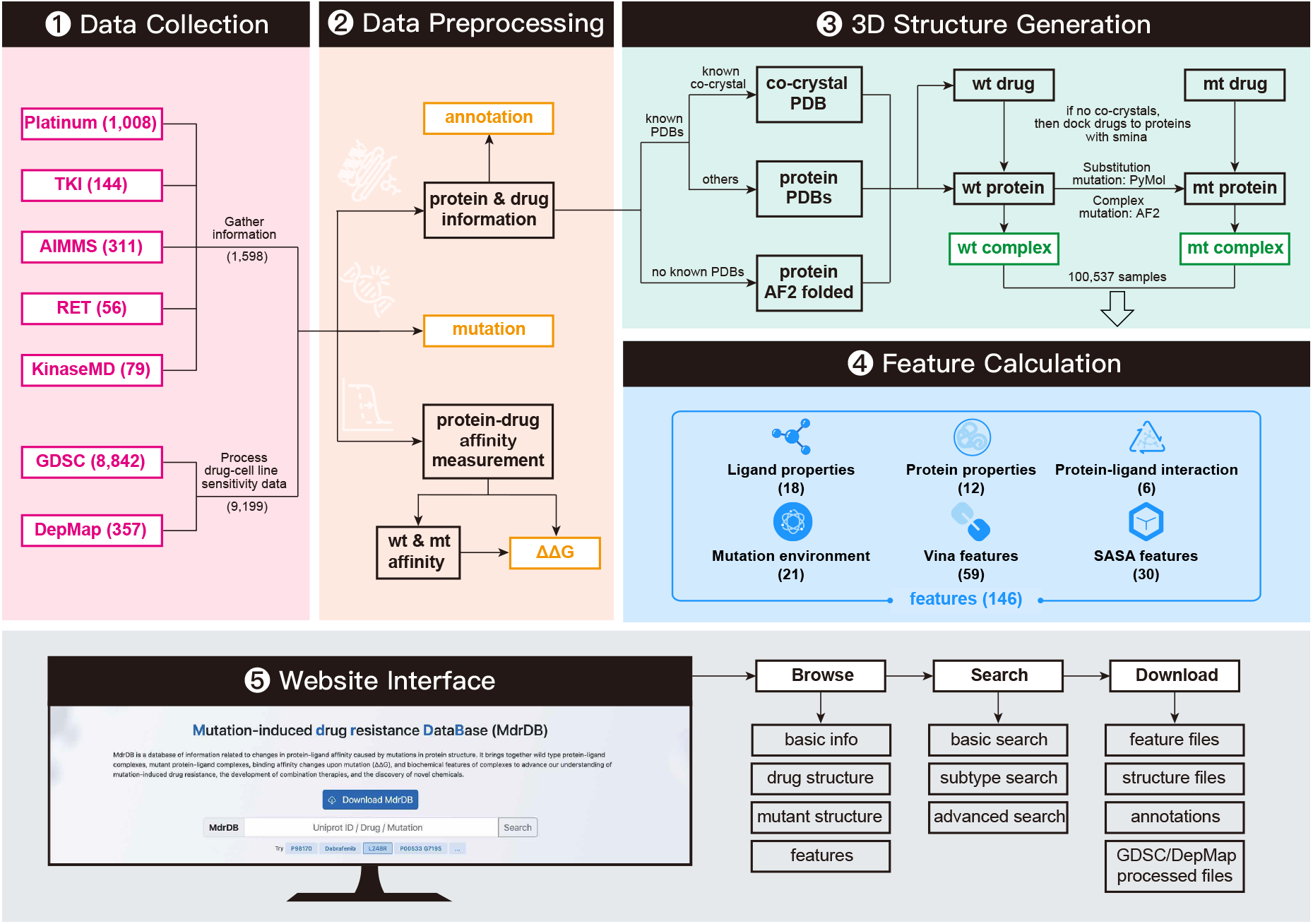
The MdrDB database construction pipeline: 1) Data collection from public data sets; 2) Data preprocessing to extract sample information; 3) 3D structure generation of protein-drug complexes for both wild type and mutant proteins; 4) Feature calculation; 5) Website construction for browsing, search and downloading of the data. Color-coded boxes: (Red) The collected publicly available datasets, the number of original samples is shown; (Yellow) Annotations, mutation information and ΔΔG values, provided in MdrDB in .tsv file format; (Green) Structural information provided in MdrDB.

### 2.1 Data Collection

MdrDB contains data from seven publicly available datasets: Platinum [17], AIMMS (Auto In Silico Macromolecular Mutation Scanning) [19], TKI [18], RET [20], KinaseMD (kinase mutations and drug response) [21], GDSC (Genomics of Drug Sensitivity in Cancer) [22], and DepMap (Cancer Dependency Map) [23]. These datasets differ slightly in terms of the information they contain.

- Platinum: provides data on affinity changes upon site mutation (ΔΔG) and experimental co-crystal structures of protein-drug complexes from the RCSB Protein Data Bank (PDB). It contains more than 1000 manually curated mutations [17].
- AIMMS: a web server for predicting site mutation-induced drug resistance, and contains a dataset of 17 protein-ligand complexes involving 311 ΔΔG values and mutations [19].
- TKI: a database of tyrosine kinase inhibitor (TKIs) resistances in ABL tyrosine kinase site mutations. A total of 144 ΔΔG values are included, along with ABL-TKI complex structures from PDB [18].
- RET is a dataset specifically related to drug resistances between three drugs (cabozantinib, lenvatinib, vandetanib) and the RET kinase mutants. 56 IC50 measurements and protein-drug complex structures were reported in the research [20].
- KinaseMD: focuses on kinase mutations, and integrates information from the Cancer Cell Line Encyclopedia (CCLE) [25] and GDSC [22] databases, and provides various annotations of drug responses on kinase mutants. 79 ΔΔG values were collected from this database [21].
- GDSC: the largest public resource for information on drug sensitivity in cancer cells. The database contains 4.4 million drug sensitivity (IC50) values across 518 drugs and 1000 cancer cell lines. It also records information related to basal expression, mutation, copy number variation, and gene methylation in the cell lines.
- DepMap: similar to GDSC, and contains drug sensitivity data on cell lines from the Achilles [26] and CCLE projects [27].

For Platinum, AIMMS, TKI, RET and KinaseMD, mutation information and mutation induced binding affinity changes are directly obtainable from the datasets. However, for GDSC and DepMap, extra steps are required to obtain the mutations and ΔΔG values (see next section).

### 2.2 Data Preprocessing

#### 2.2.1 GDSC/DepMap raw data processing

Mutation and ΔΔG values were obtained from these two datasets via a process similar to that used by KinaseMD: (i) Mutation information acquisition; (ii) Drug-cell line response calculation; and (iii) ΔΔG calculation.

Cell lines were first grouped into wild type and mutated samples for a specific protein, and mutation information for the protein was gathered. Specifically, for a particular protein, cell lines that did not have mutations on them were considered to be control (wild type samples), while cell lines with mutations were considered to be mutant. Mutations were then filtered to retain only six types of mutation: single substitution, multiple substitution, deletion, insertion, indel and complex mutations (i.e., combinations of the other five mutation types).

Then, we filtered the drugs with known protein targets and queried drug sensitivity (IC50 values) data on all cell lines. For drugs with multiple known protein targets, each protein was considered individually. If, in the cell line, only one of these proteins was mutated, the cell line was kept. Otherwise, the cell line was discarded. After this, for each (protein, cell line, drug) mutant sample, the mutation string was generated by merging all mutations of the mutant protein; IC50 values were averaged over all cell lines, and this average value was taken as the mutant protein-drug affinity value IC50(MT). A corresponding wild type sample was similarly assigned and IC50s again averaged over cell lines, and this average value taken as the wild type protein-drug affinity IC50(WT).

Finally, these affinity pairs were used to calculate ΔΔG according to the formula [28]:

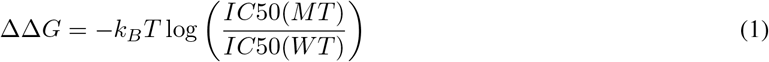

#### 2.2.2 Sample data consolidation

From all seven datasets, five basic pieces of information were collected and consolidated: protein name, uniProt ID, mutation string, drug name, and ΔΔG value. UniProt IDs were identified by querying UniProtKB [29] with protein names. The consolidated data was then divided into separate tables, with samples of the same mutation type kept together. For samples from Platinum and TKI, wild type and mutant protein-drug complex structures were also obtained from the original datasets.

#### 2.2.3 Protein and drug annotations

For protein annotations, we used the Interpro API [30] to query the protein family, homologous superfamily and domain information. For drug annotations, we used the PubChem PUG REST API [31] to query the CID, Depositor-Supplied Synonyms, FDA machanism, MeSH (Medical Subject Headings) and Drug Classes information.

### 2.3 Three-Dimensional Structure Generation

With the exception of samples from Platinum and TKI which already directly include 3D structure information, we prepared 3D structures of wild type proteins, mutant proteins and drug binding poses for all samples from the other datasets.

#### 2.3.1 Wild type protein structure preparation

For each protein (UniProt ID), all associated PDBs were identified with the RCSB REST API [32]. The .pdb or .cif for 3D structures and .fasta for sequences were downloaded from RCSB PDB. Then, all water, solvent, and ions were removed from the PDB files. Next, the protein chains and ligands were split into separate files. Each chain was annotated and only the chains corresponding to the UniProt ID were kept. If multiple chains were found to exist for protein, the longest one was kept. Meanwhile, the largest ligand was kept as the docking box generation reference. Finally, each mutation for a protein was checked against all available PDBs. If the mutation sites could be found in the PDB, the mutation and drug would be assigned to the PDB.

#### 2.3.2 Mutant protein structure generation

We generated mutant structures from wild type structures using PyMOL [33] or from wild type sequences using AlphaFold 2.0 [24]. Specifically, the Mutagenesis Wizard module of pymol-open-source v2.5.0 was used to generate a mutation by replacing a residue with a new amino acid type, sample several rotamers from the rotamer library and generate several non-clashing conformations. Then, the most likely rotamer was chosen as the mutated residue. For AlphaFold 2.0, we used the protein amino acid sequence as the input to predict the structures. A length threshold of 2,000 was set for computing resource considerations. For MSA (Multiple Sequence Alignment) searching, ‘reduced_db’ was used. For inference, the ‘monomer_ptm’ models were used. Five models were generated and the one with highest averaged pLDDT (predicted lDDT-Cα) value was chosen as the predicted structure for further procedures. Several rules were followed to decide which tool was used for mutant structure generation. For proteins with known PDBs containing the mutation sites:

- For single substitution and multiple substitution mutations, PyMOL was used for mutant protein generation.
- For deletion, insertion, indel and complex mutations, AlphaFold 2.0 was used for mutant protein structure prediction. For fair comparison and feature calculation, post-processing was carried out to keep the residue numbers the same except at the mutated sites.

For proteins with no known PDBs containing the mutation sites, AlphaFold 2.0 was used for both wild type protein and mutant protein structure prediction. Structures with an average pLDDT larger than 70 for the whole structure were kept, which was a confidence threshold for the predicted structures in AlphaFold 2.0. In addition, if a mutated site was located on a region that was poorly predicted (pLDDT < 50), the sample was also discarded. After structure generation, the mutant protein was aligned with the wild type protein. This alignment was carried out by only taking the backbone atoms of both wild type and mutant residues whose pLDDTs were larger than 70 into consideration.

#### 2.3.3 Drug structure generation

SMILES strings for all drugs were first obtained using the PubChem PUG REST API [31] (several drugs that could not be directly identified via PubChem were manually checked and assigned). Ions and salts were then removed from the SMILES, the structures neutralized, and the resulting SMILES strings rewritten in canonical format. 3D structures were first generated using Open Babel 3.1.1 [34], using the ‘–gen3D’ flag, and non-polar hydrogens added to the generated structures with the ‘–addpolarH’ flag. After structure generation, molecular docking was carried out with smina [35], using default docking parameters:

- If the wild type PDB contained a known in-pocket ligand, then ‘–autobox_ligand’ was selected.
- If no ligand was present, the whole protein was used to generate the docking box.

After docking, the conformation with the best smina score was kept.

### 2.4 Feature Calculations

Based on the structures of the wild type and mutant protein-ligand complexes, we calculated a total of 146 biochemical features relevant to machine learning prediction of mutation-induced affinity changes. These features were first used in the work of Aldeghi et al. [36], and we follow the procedures in their original paper for their calculation:

- 18 ligand properties such as logP, molecular weight, number of hydrogen acceptors and donors were calculated using RDKit.
- 12 features describing the changes for the mutated amino acids were calculated (again with RDKit), such as hydrophobicity, number of heavy atoms, and the change in side-chain volume.
- 21 features describing the mutation environment were calculated with Biopython, including the distribution of ligand and protein atoms around the mutation site and the number of residues in different property groups.
- 6 features related to the protein-ligand interactions were calculated using the Protein-Ligand Interaction Profiler (PLIP) [37]: hydrogen bonds, hydrophobic contacts, salt bridges, π-stacking, cation-π interactions, and halogen bonds.
- The Vina binding score as well as 58 scoring function terms were calculated using Delta-Vina XGBoost [38, 39].
- 30 pharmacophore-based solvent accessible surface area (SASA) features are calculated using Delta-Vina XGBoost. [39]

With the exception of the ligand property features, the numerical values *x* of all other features were given in MdrDB correspond to the difference between the mutant (*x_mt_*) and wild type values (*x_wt_*), i.e., *x* = *x_mt_* - *x_wt_*.

## 3 DATABASE CONTENT AND USAGE

Table 1 summaries the data contained in MdrDB. A user-friendly website (https://quantum.tencent.com/mdrdb/) provides access to the curated data and structure information on wild type and mutant complexes, and provides functions for browsing, searching, displaying and downloading the data.

**Table 1:** Overview of data represented in MdrDB.

| Property | # counts | # counts in source datasets |  |  |  |  |
| --- | --- | --- | --- | --- | --- | --- |
| # Samples | 100,537 | AIMMS (5,048), DepMap (1,500), GDSC (91,215), KinaseMD (1,334), Platinum (840), RET (456), TKI (144) |  |  |  |  |
| # Unique UniProt IDs | 240 | AIMMS (10), DepMap (5), GDSC (99), KinaseMD (31), Platinum (126), RET (1), TKI (4) |  |  |  |  |
| # Unique ligands | 440 | AIMMS (16), DepMap (5), GDSC (180), KinaseMD (48), Platinum (126), RET (3), TKI (8) |  |  |  |  |
| # Mutations | 2,503 | AIMMS (121), DepMap (49), GDSC (1,805), KinaseMD (43), Platinum (530), RET (11), TKI (31) |  |  |  |  |
| # Unique Uniprot ID-Mutations | 2,673 | AIMMS (173), DepMap (49), GDSC (1,818), KinaseMD (43), Platinum (578), RET (11), TKI (80) |  |  |  |  |
| # Unique PDB IDs | 5,119 | AIMMS (430), DepMap (220), GDSC (4,128), KinaseMD (802), Platinum (241), RET (21), TKI (8) |  |  |  |  |
| Mutant structure source | # Platinum processed | # TKI processed | # PyMOL mutated | # AlphaFold2 folded |  |  |
|  | 840 | 144 | 94,987 | 4,566 |  |  |
| Drug pose source | # Platinum cocrystal | # TKI cocrystal | # TKI docked | # Docked |  |  |
|  | 840 | 131 | 13 | 99,533 |  |  |
| Mutation types | # Single substitution | # Multiple substitution | # Deletion | # Insertion | # Indel | # Complex |
|  | 84,038 | 11,977 | 3,228 | 355 | 243 | 696 |

### 3.1 Data summary

#### 3.1.1 Samples

At the time of writing, MdrDB contains 100537 samples, generated from 240 proteins (5119 total PDB structures), 2503 mutations, and 440 drugs. 95971 of the total samples are based on available PDB structures, with the remaining 4566 samples predicted with AlphaFold 2.0.

#### 3.1.2 Mutation types

In addition to single-point substitution mutations and multiple-points substitution mutations, MdrDB also contains complex mutations including deletion, insertion, and indel (insertion-deletion) mutations, as well as multiple-site mutations containing a number of the aforementioned mutations. Figure 2A shows MdrDB samples categorized by their mutation types. Approximately 83.6% of the samples are single substitution mutations; 11.9% are multiple substitution mutations; and 4.5% are complex mutations, of which deletion mutations account for the largest proportion.

**Figure 2:**
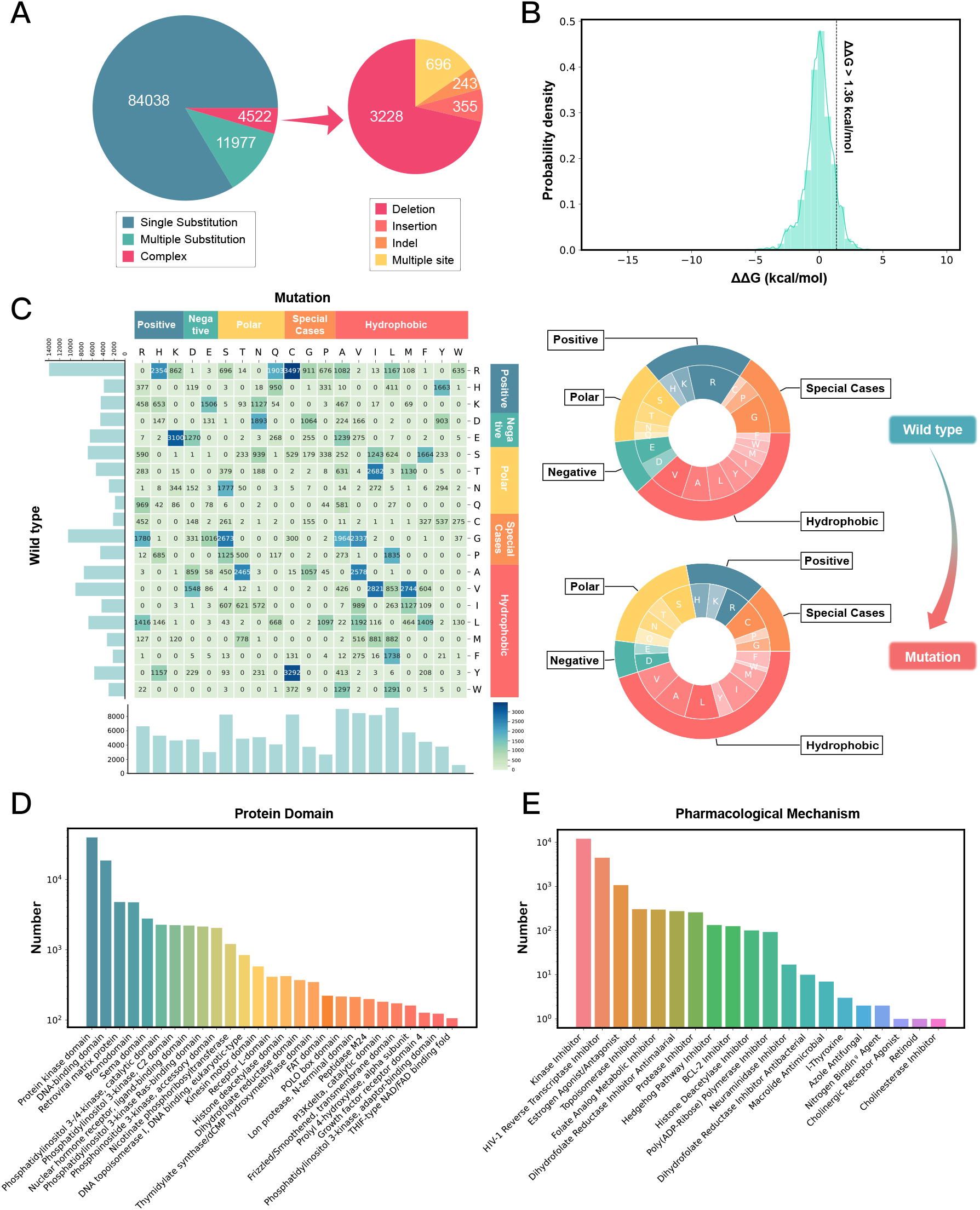
MdrDB mutation statistics, ΔΔG distribution, and protein and drug annotations. (A) Number of samples of each mutation type. ‘Others’ includes four mutation types: deletion, insertion, indel, and complex. (B) Histogram of protein mutation-induced ligand binding affinity changes measured by ΔΔG (kcal/mol). Line at ΔΔG = 1.36 kcal/mol separates mutations defined as resistant from susceptible. (C) Number of amino acid changes from substitution mutations. (Left) Heat map shows the number of amino acid changes from substitution mutations, with different colors corresponding to the number of changes in different amino acids. The number of samples for each amino acid in the wild type (mutation) is displayed as a bar chart along the left axis (bottom) of the plot. (Right) Donut charts show the number of amino acids in the samples corresponding to substitution mutations. The proportion of each amino acid in wild-type (mutated) samples is displayed top (bottom). (D) Number of samples annotated by protein domains. X-axis gives protein domain, with y-axis the corresponding sample number on logarithmic scale. (E) Number of samples annotated by pharmacological mechanisms.

#### 3.1.3 Mutation-induced drug resistance

The distribution of mutation-induced ligand binding affinity changes, measured by ΔΔ*G*, is shown in Figure 2B. A standard for defining resistant samples was given in [18], where samples with a larger than 10-fold drop in binding affinity, corresponding to ΔΔ*G* > 1.36 kcal/mol, are considered to be resistant. Based on this standard, 8197 (8.2%) samples in MdrDB are mutation resistant.

We analyzed the distribution of amino acid types in substitution mutation samples, before and after mutation, to shed light on mutation preferences. With the 20 natural amino acids divided into five groups – positively charged, negatively charged, polar, hydrophobic, and three amino acids classified as special cases (Figure 2C and Figure S1A) – we find the following: Overall, 30.8% mutations are observed within the same amino acid group, while 69.2% mutations occur across different amino acid groups. Arginine (R) is the most frequently mutated amino acids, followed by glycine (G) and valine (V). Leucine (L), alanine (A), serine (S) and cystenine (C) are the most frequently mutated amino acids. Meanwhile, the amino acids in charged groups do not show a marked overall bias for mutation. However, different amino acids within charged groups do display preferences for mutation. For example, histidine (H) prefers to mutate to tyrosine (Y, 42.8%), while glutamic acid (E) prefers to mutate to lysine (K, 48.2%). In contrast, amino acids in the polar, hydrophobic and special groups have a higher preference to mutate to amino acids in hydrophobic groups, with ratios of 57.1%, 56.0%, and 44.3% respectively. Similar conclusions holds for the MdrDB CoreSet (Figure S1B and Figure S2), where non-repetitive samples considering only unique ‘protein-drug-mutations’ are kept.

In addition, we analyzed the secondary structure elements where mutations occurred. As shown in Figure S4, the ΔΔ*G* value distributions were not significantly different across samples where mutations are located in *α*-helix, *β*-sheet or loops.

#### 3.1.4 Protein and drug annotation

Annotated for the proteins and drugs are provided for each sample in MdrDB. For proteins, a total of 160 kinds of protein domains, 154 protein families and 143 protein superfamilies can be found in MdrDB (see Supplementary for more information). The protein domains with at least 100 samples are shown in Figure 2D. Protein kinase domains account for the largest proportion (39.5%) of all samples, followed by DNA-binding domains (18.5%), retroviral matrix proteins (4.8%) and bromodomains (4.7%). For drug mechanism annotations, a total of 20 FDA documented pharmacological classes, 60 MeSH pharmacological classes and 9 PubChem drug classes can be found in MdrDB (see Supplementary for more information). 67 (15.2%) out of all 440 drugs have known FDA pharmacological classes, and the corresponding samples were counted and shown in Figure 2E. The largest proportion of samples take kinase inhibitors as drugs (12.0%), followed by HIV-1 reverse transcriptase inhibitors (4.5%).

### 3.2 Web design and interface

The web interface of MdrDB was implemented using caddy2 and Bootstrap v5.0. All processed data, structure files, biochemical features and annotations were stored in Tencent Cloud Object Storage (COS). Interactive charts were implemented using Apache ECharts. A full tutorial for MdrDB is available at https://quantum.tencent.com/mdrdb/tutorial.

#### 3.2.1 Browse

Users can browse all the curated and processed data through the Search MdrDB page, accessible via a quick-access button in the navigation bar. Data is presented in tabular form, with the following basic data given for each sample: MdrDB sample ID, UniProt ID, PDB ID, mutation type, mutation string, drug name, drug SMILES, ΔΔG value, and sample source. From this page, users can easily search, sort, filter and download samples (Figure 3A).

**Figure 3:**
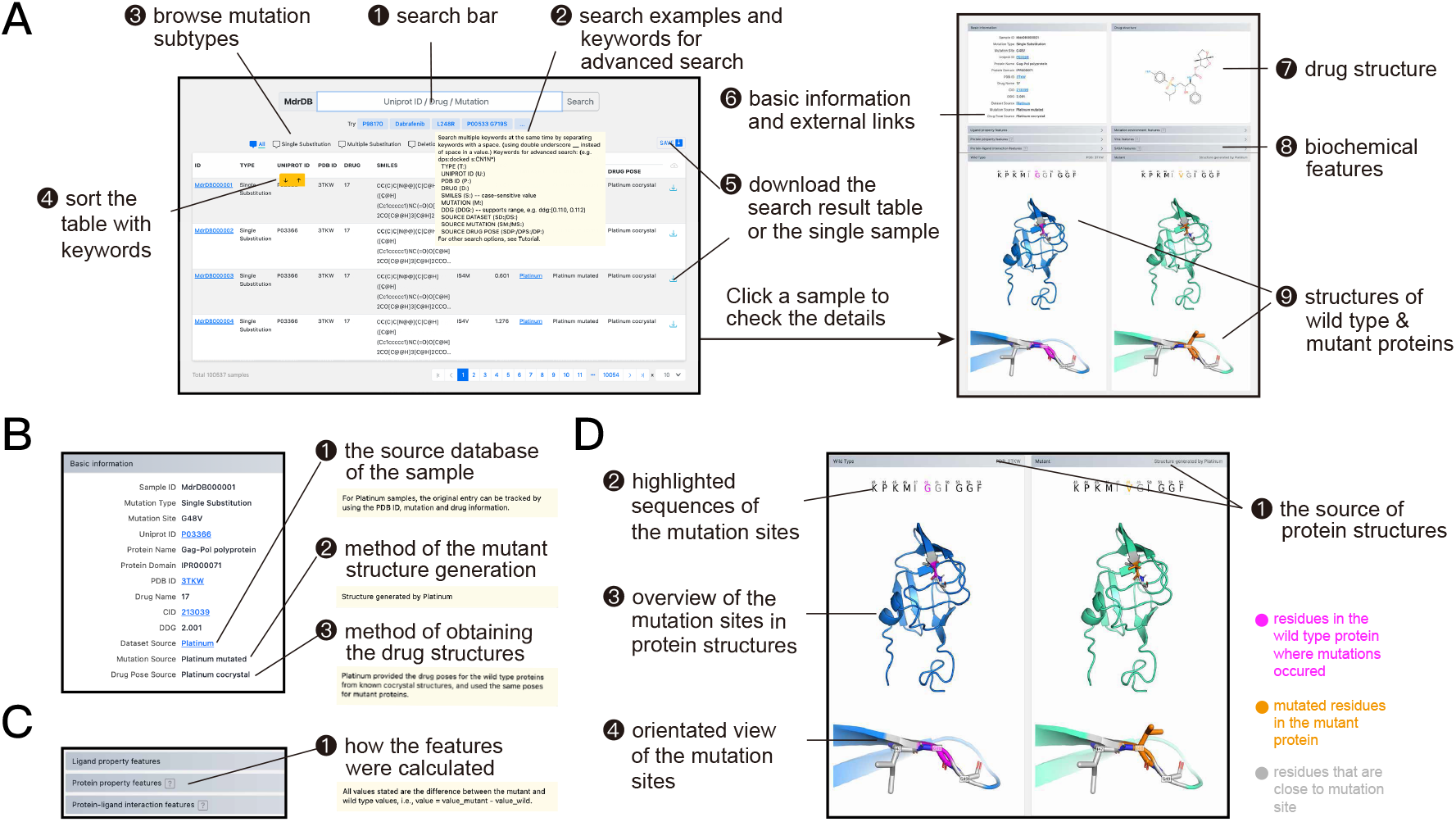
The search and browse page and the sample display page of the MdrDB website. (A) Basic functions of the search and browse page and information included in the sample display page. (B) Details of data tracking in the “Basic Information” section on the sample display page. (C) Details of feature calculation in the “Features” section on the sample display page. (D) Details of different views of wild type and mutant structures in the “Protein Structures” section on the sample display page.

#### 3.2.2 Search

A full guide to the basic search and advanced search provided by MdrDB is contained in the tutorial available from the website. Here we summarize the key features.

With basic search, users can query any of the following keywords directly in the search bar (Figure 3A.❶):

- MdrDB ID: e.g., “MdrDB00158”
- UnitProt ID: e.g., “P00533”
- Mutation String: e.g., “G791S”
- Drug Name: e.g., “Embelin” (alternative names of drugs recorded in PubChem may also be used)

Advanced search (Figure 3A.❷) is performed by prepending queries to keywords with an appropriate prefix, e.g., “MS:” corresponds to the MUTATION_SOURCE keyword. An example of a valid query is “MS:alphafold2”.

The “*” wildcard symbol can be used to search in place of unspecified characters or values, e.g., “P:*G9*” or “T:*substitution”.

Multiple (space-separated) keywords can be queried simultaneously with both basic and advanced search. For example, UniProt ID and mutation string keywords can be searched simultaneously, with a query such as “P98170 R443C”.

#### 3.2.3 Display

On the sample display page, users can view detailed sample information (Figure 3A❻-❾): (i) basic information, (ii) drug structure, (iii) biochemical features, and (iv) structure of wild type and mutant proteins. The basic information block (Figure 3B), in addition to containing the information displayed in the browse and search page, also gives hyperlinks to the corresponding UniProt (UniProt ID), RCSB PDB (PDB ID) and PubChem (CID) entries. The drug structure block (Figure 3A.❼) displays the two-dimensional (2D) structure of the drug. The biochemical features block (Figure 3A.❽) contains six types of features: ligand property features, protein property features, mutation environment features, protein-ligand interaction features, Vina features, and SASA features. The structure of wild type and mutation block (Figure 3D.❷) displays the sequence around the mutation sites, overall structures of wild type and mutant proteins, and close-up views of the mutated sites. Color coding is used to indicate mutation site residues in the wild type (magenta) and mutant (orange) proteins. The two residues that are next to the mutation site are colored grey. The close-up view gives a clear picture of the changes in residues near the mutation site before and after the mutation. Users can download structure and feature files for each samples by clicking the download button on the upper right of the sample display page.

#### 3.2.4 Download of data, figures, and tables

All plots displayed on the website (e.g., pie charts, bar plots and heatmap) can be downloaded in a variety of formats (JPG, PNG, and PDF) by clicking the button at the top right of the plot. On the download page of the MdrDB website, users can download metadata, structure files, processed biochemical features and annotation files. MdrDB provides two dataset types to users for downloading: MdrDB_CoreSet and MdrDB_FullSet. MdrDB_CoreSet provides non-repetitive “UniProt-mutation-drug” samples whose features are averaged over all corresponding PDB features. MdrDB_FullSet provides all “UniProt-mutation-drug” samples, whose features are calculated based on each PDB structure.

## 4 CONCLUSION

Here we have introduced MdrDB, the largest drug resistance database to provide all information highly relevant to protein mutation-induced drug binding affinity changes. In addition to basic information regarding the proteins, drugs, mutations and changed affinity, and structural data for wild type and mutant complexes, 146 calculated biochemical features and extra annotations are also provided. These features and annotations were chosen to be convenient to use for model training and sample splitting in machine learning algorithms for drug resistance prediction [40]. In addition, MdrDB is also the first database that includes drug resistance related mutation types beyond substitution mutations, and the availability of wider and more complex mutation types can be used to test the generalization of machine learning models.

A comprehensive resource providing a variety of information for studying protein mutation, predicting drug resistance and discovering novel chemical compounds, MdrDB will be updated regularly and include more public data in the future.

## 5 DATA AVAILABILITY

All data is available to browse and download on the MdrDB website (https://quantum.tencent.com/mdrdb/).

## Notes

https://quantum.tencent.com/mdrdb/

### Competing Interest Statement

The authors have declared no competing interest.

